# Influence of internal seiche dynamics on vertical movement of fish

**DOI:** 10.1101/2021.08.03.454964

**Authors:** Ivan Jarić, Milan Říha, Allan T. Souza, Rubén Rabaneda-Bueno, Vilem Děd, Karl Ø. Gjelland, Henrik Baktoft, Martin Čech, Petr Blabolil, Michaela Holubová, Tomáš Jůza, Milan Muška, Zuzana Sajdlová, Marek Šmejkal, Lukáš Vejřík, Ivana Vejříková, Jiří Peterka

## Abstract

Internal seiches are common in stratified lakes, with significant effects on stratification patterns, hydrodynamics and vertical nutrient transport. In particular, seiche can change the vertical distribution of the thermocline and the cold hypolimnetic and warm epilimnetic water masses by several meters on a timescale of a few hours. The results are rapid and strong changes in temperature profiles and oxygen availability that can have profound effects on vagrant and sessile organisms. Internal seiche dynamics could therefore affect fish communities directly through physiological stress and elevated mortality, and indirectly through prey distribution.
The aim of this study was to analyze the effects of internal seiche dynamics on lacustrine fish behaviour, and to characterize fish reaction patterns, with the main focus on vertical movement of fish in the vicinity of a shifting thermocline, and avoidance of cold hypolimnetic water.
The analysis was based on acoustic telemetry data from Lake Milada, a post-mining lake in the Czech Republic, with a total of 55 tracked individuals of four species: northern pike (*Esox luciu*s), wels catfish (*Silurus glani*s), tench (*Tinca tinc*a) and rudd (*Scardinius erythropthalmu*s).
The effects of seiche dynamics on the four species studied were weak but significant during the day, but only on rudd during the night. Upward seiche produced stronger reactions in fish than downward seiche, and the effects were manifested only during the strongest seiche events.
Thermocline shifting during seiche events may induce a transient reduction in habitat for seiche-reacting species, thus potentially affecting predation and other inter- and intra-specific interactions, and probably affecting fish community dynamics.

**Graphical abstract:** 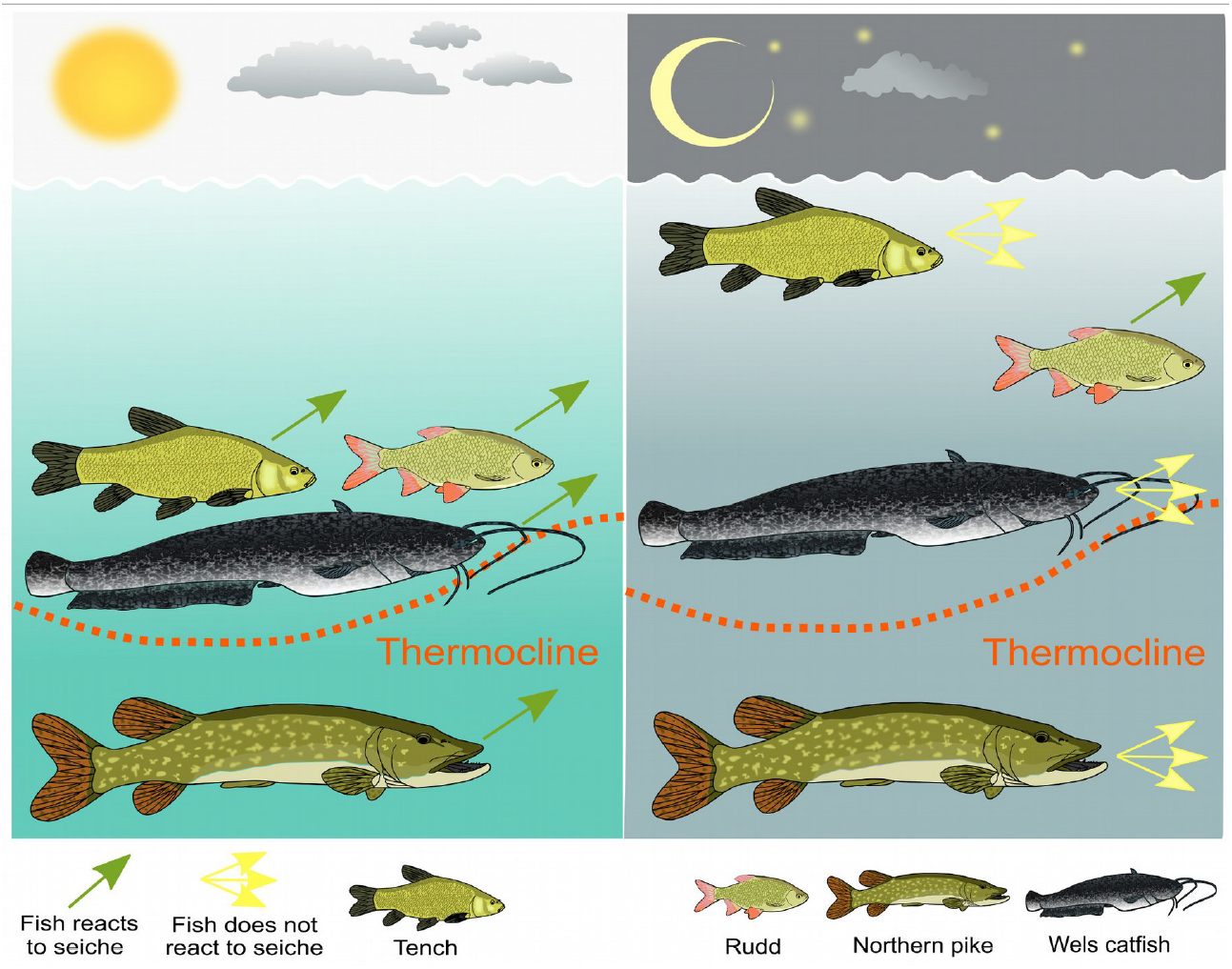

## 1. Introduction

Water temperature is a critical environmental factor for aquatic ectotherms (Cossu et al., 2017). Such species are typically adapted to limited temperature ranges, with distinct, speciesspecific thermal optima at which their physiological performance is maximized (Aspillaga et al., 2017). Rapid temperature changes can induce thermal stress with a range of physiological and behavioural responses, potentially leading to death in extreme cases (Donaldson et al., 2008). Nonterritorial fish move to more suitable habitats, among other factors, to adjust their surrounding temperature exposure. This can lead to thermally stratified fish community distribution that can be long-lasting or transient, depending on the duration of the thermal stratification (Coulter et al., 2016).

Temperature fluctuations are frequent in the vicinity of a thermocline, and are especially relevant in temperate lakes (Cossu et al., 2017). Internal seiches, or lacustrine internal waves, represent a common hydrological phenomenon that causes large fluctuations of vertical temperature stratification by changing thermocline depth, normally as a result of wind-generated pressure gradients (Ostrovsky et al., 1996; Stevens & Lawrence, 1997). They are common in stratified water bodies such as lakes, but are ephemeral and often go undetected (Emery, 1970). Internal seiche affects water stratification, hydrodynamics, vertical nutrient transport, temporary temperature profiles and oxygen availability. As such, internal seiche dynamics has the ability to influence fish communities (Stevens & Lawrence, 1997), and lead to changes in vertical distribution of fish communities. Its impact is also manifested indirectly, through changes in the vertical distribution of prey, predator and competitor species. As the wind patterns are currently changing around the globe due to climate change (Seneviratne et al., 2012), seiche events are also likely to intensify in frequency and strength over time (Kirillin & Shatwell, 2016), which makes it important to understand the consequences of the phenomenon on fish communities.

However, except for a couple of case studies (Levy et al., 1991; Easton & Gophen, 2003), internal seiche effects on fish distribution have not received proper attention, and remain largely unknown. The main reason for such a lack of studies is probably that internal seiches are relatively rapid and ephemeral events that affect only a narrow range of the water column (usually no more than a few meters), making detection of fish movement and distribution complicated using conventional fish sampling methods. Recent developments in advanced research techniques, such as autonomous telemetry positioning systems (Baktoft et al., 2015), offer the opportunity to address this question at a level of detail previously not possible.

In the present study, we examined behavioural and distributional responses of four fish species to internal seiche dynamics in a natural lake environment using high-resolution spatiotemporal telemetry tracking. Species selected for the study are reported to differ in temperature preferences and life strategies. The study was focused on two top predator species, northern pike (*Esox lucius*) and wels catfish (*Silurus glanis*), and two generalist and potential prey species, rudd (*Scardinius erythrophalmu*s) and tench (*Tinca tinca*). Northern pike is described as a cool water species, with a wide range of preferred temperatures (Nordahl et al., 2020; Pierce et al., 2013). Similarly, rudd is considered to be temperature-tolerant species (García-Berthou & Moreno-Amich, 2000; Hicks, 2003; Kottelat & Freyhof, 2007), while wels catfish and tench are documented as species preferring warmer water (Capra et al., 2018; Cucherousset et al., 2018). Species-specific temperature preferences can be assumed to affect response to seiche dynamics, and a lower intensity of responses to seiche dynamics can be expected for temperature-tolerant species than for warmwater species.

Fish reaction to seiche dynamics can also be dependent on a diel period. Regular diel alteration of vertical distribution is a common phenomenon in aquatic environment, and it was also documented for studied species (e.g. Bohl, 1979; Mehner, 2012; Nordahl et al., 2020; Slavík & Horký, 2012). Fish tend to be closer to the surface during nighttime, but opposite patterns and individual differences have also been documented (e.g., for pike; Nordahl et al. 2020). As such, fish might be exposed to internal seiche dynamics only during one diel period.

Lastly, individual variation can represent an important driver of behavioral responses to seiche dynamics. High level of individual variation was observed in behavior of many animal taxa (Bell et al., 2009; Dingemanse & Wolf, 2013), and individual differences in movement behavior were found in northern pike (Nyqvist et al., 2018; Říha et al. 2021), wels cactfish (Slavík & Horký, 2012) and different cyprinid species (e.g. Hulthén et al., 2017; Říha et al., 2015; Winter et al., 2021).

The aim of this study was to analyze fish behaviour and vertical movement in the vicinity of a shifting thermocline, as well as inter- and intra-specific differences in reaction patterns. We established the following hypotheses: 1) fish do react to internal seiche dynamics by changing their vertical position, 2) reaction patterns to internal seiche dynamics will differ among species based on their ecology and life history, 3) they will differ between day and night, and 4) they will also vary strongly among individuals.

## 2. Materials and methods

The analysis of internal seiche dynamics and its effects on fish behavior was based on a fish telemetry dataset from the Lake Milada (50°39′N, 13°58′E) in Czech Republic (Fig. 1), a 250 ha large water body created after aquatic restoration of a mining pit during 2001-2010 (Vejřík et al., 2017). It is an oligotrophic lake, with the mean and maximum depth of 16 and 25 m, respectively, and with poorly oxygenated water below 15 m depth. The lake has a simple, elongated shape, with a flat, U-shape bottom profile, and an open surrounding landscape, with no forest cover or other structures that could interrupt or weaken wind flow. It has a suitable position for studying internal seiche events, as the prevailing winds tended to follow the direction that closely follows the longer axis of the lake, and allows frequent occurrence of internal waves (Čech et al., 2011; Fig. 1).

**Figure 1.**
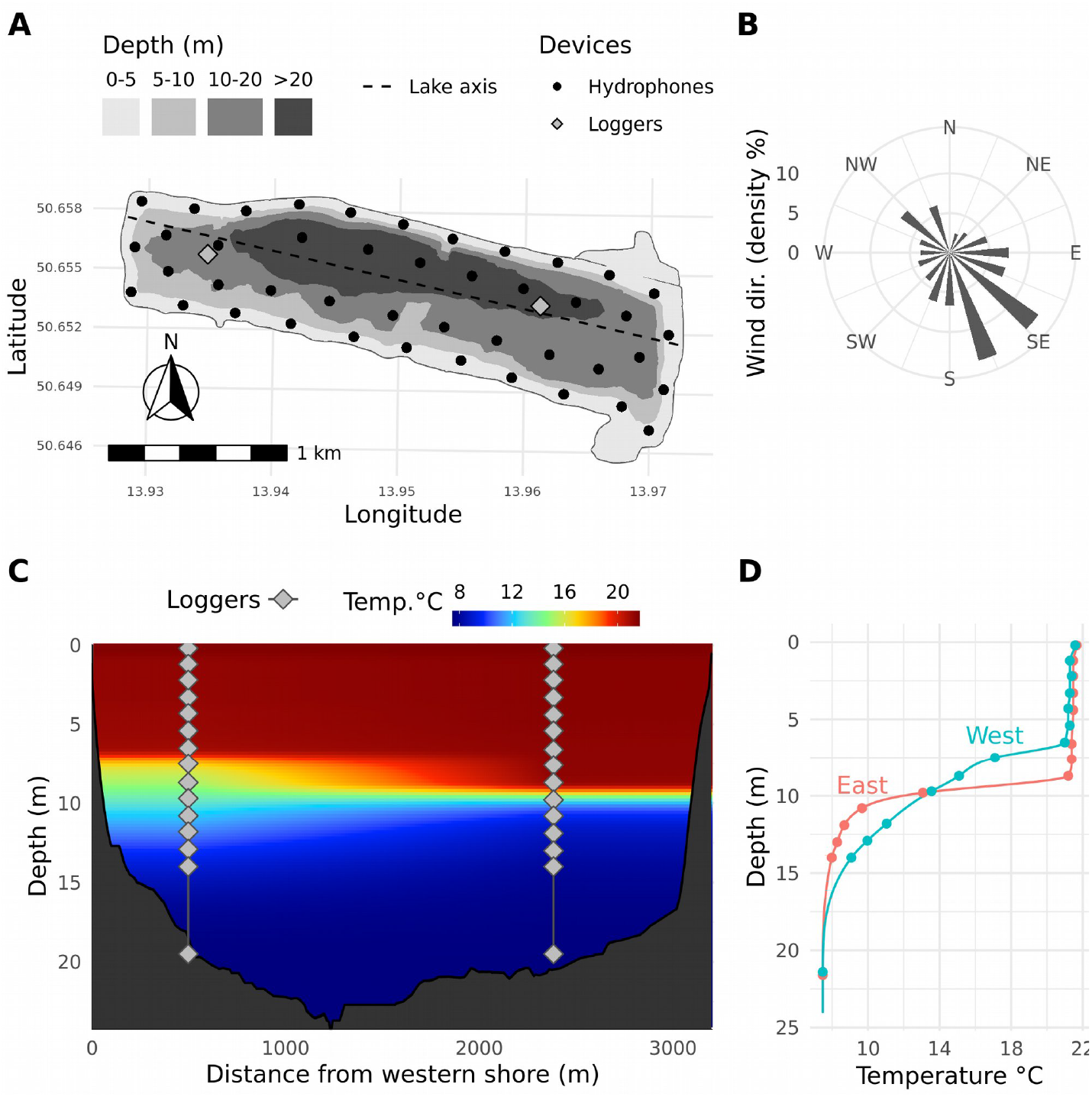
Outline of the studied lake (Milada Lake, Czech Republic), with isobaths and the positions of acoustic receivers and two HOBO dataloggers (A), wind direction during the studied period (B), vertical positions of loggers and temperature distribution during one seiche event (C), and measured and interpolated temperatures at the both logger sites during the same seiche event (D).

Between April 2015 and March 2016, a positioning system array with 44 receivers (Lotek Wireless Inc., Canada, model WHS3250) was deployed in the lake, with a distance of 203 to 288 m between each of the three closest receivers (mean 251 ± 18 m) and a depth range of 4.5 to 5.5 m. Exact receiver positions were measured with a differential GPS-unit (Trimble, USA). Testing of the system, performed prior to the study (September 2014), indicated full lake coverage, while system accuracy was ensured by 10 stationary reference tags (Lotek Wireless Inc., Canada, model MM-M-16-50-TP, burst rate 25 secs) placed at different sites and depths throughout the lake (Fig. 1). Additional tests of accuracy were conducted both after deployment of the system and before retrieval, by dragging the reference tags by a boat.

Fish were captured between 5 and 10 May 2015 by electrofishing, longlining and angling (Vejřík et al., 2019). After capture, fish were anaesthetized with 2-phenoxy-ethanol (SIGMA Chemical Co., USA; see supporting information Supp1), measured, weighed and tagged. Tagging was performed by making a 1-1.5-cm-long incision on the ventral surface, inserting the transmitter (Lotek Wireless Inc., Canada, with pressure and motion sensors, and burst rate 25 s; see supporting information Supp1) into the body cavity, and closing the incision using two separate sutures (see supporting information Supp1 for information on tagging procedure). All fish were released immediately after recovery from anaesthesia.

The fish telemetry dataset comprised high-resolution spatiotemporal data on the vertical and horizontal positions of 55 individuals of four fish species: northern pike, wels catfish, tench, and rudd. All captured fish were large adults (Table 1).

**Table 1.** Number of specimens and the total (TL) and standard (SL) body length and weight (W) of the individuals tracked with acoustic telemetry in Milada Lake (mean ± standard deviation).

|  | N | TL (cm) | SL (cm) | W (kg) |
| --- | --- | --- | --- | --- |
| Rudd | 14 | 385.4 ± 23.2 | 327.1 ± 25.6 | 1.08 ± 0.17 |
| Tench | 16 | 486.3 ± 33.0 | 413.4 ± 29.7 | 2.21 ± 0.42 |
| Northern pike | 11 | 785.9 ± 142.2 | 694.5 ± 131.3 | 3.72 ± 2.08 |
| Wels catfish | 14 | 1178.6 ± 222.2 | 1090.0 ± 208.2 | 10.72 ± 6.02 |

Transmitter detection data were processed with User Managed Acoustic Positioning (U-MAP, Lotek Inc.) software, which estimates transmitter positions through multilateration of time-of-arrival data detected at three or more receivers. The algorithm solves nonlinear equations with multiple solutions, which are consequently exported for postprocessing to eliminate erroneous position estimates. The performance of position estimates and possible filters was evaluated by: 1) short-term tag-tows from a boat with a high-precision GPS-device, 2) several stationary reference tags placed in the lake throughout the study, and 3) visual inspection of fish track data. Detailed information on data filtering is provided in supporting information Supp2 (see also Říha et al. 2021).

Identification of thermocline and internal seiches was based on the two sets of dataloggers (Onset, USA, HOBO Pendant temp/light 64K), placed in the eastern and western section of the lake during the experiment, with 1890 m distance (Fig. 1). At each of these two sites, 14 dataloggers were spread from the surface to 13 m depth (1 m distance), with an additional datalogger placed at 20 m depth. As such, the dataloggers completely covered the depths at which the thermocline is found throughout the year, as well as the area where the internal seiche zone may be detected. Dataloggers measured temperature at 5-minute intervals.

Temperature profiles from each logger site and each 5-minute interval were smoothed by *monotone* cubic spline using Hyman filtering and mid-thermocline was defined as depth with greatest density gradient. Additionally, the depth of the top and base of the thermocline were calculated as depth range limits with a density gradient greater than 0.2 kg/m^3^ per meter. Computations were performed using R package *rLakeAnalyzer* (Read et al., 2019; see also Read et al., 2011), specifically, using the functions *thermo.depth* and *meta.depths*. In special cases, such as periods with multiple thermoclines or with a global temperature profile density gradient close to threshold, the results yielded a significant amount of noise that was removed by temporal smoothing. First, isotherms were computed using the moving median of thermocline temperatures within a three-day window. To obtain final thermocline depth, the isotherms were smoothed by Fast Fourier Transform smoother using only the signal frequencies corresponding to the wave period of 4h30 and longer.

We established three metrics to characterize the thermocline in our statistical models: mean temperature gradient between the top and base of the thermocline (mean_gradient), seasonal thermocline depth (seasonal_depth) computed by applying a general additive model (GAM) smoother on thermocline depth (formula depth ~ s(time, k = 100) on n = 38800 points) and difference between the seasonal depth and the actual depth of the thermocline (amplitude), which represented the strength of the seiche.

Fish detections were assigned a temporally closest horizontal position and detections more than 15 minutes from the closest position were excluded. All positions and the two logger sites were projected onto the lake axis connecting the western and eastern shores of the lake (Fig. 1). Thermocline descriptors for each fish detection were obtained by linear interpolation of the values at each logger site. For analysis of fish reaction patterns in the presence of a seiche, we focused on periods when the difference in thermocline depth between the east and west logger sites was greater than 0.7 m (vertical resolution of fish tags).

We focused on the period between June 16 and October 15, which represents a period of a stable thermocline/water stratification. At the same time, it represents the peak of the growing season of macrophytes (Vejříková et al., 2017) and the peak of activity of studied species due to higher values of water temperature. As a validation of identified seiche dynamics, we checked concordance between wind patterns and seiche dynamics, which showed good overlap between the two (Fig. 2). Hypoxic conditions (oxygen concentrations below 2 mg O_2_/L) appeared in the lake only late in the summer (September and October) in deeper layers of the water column (below 15 m in the middle of September).

**Figure 2.**
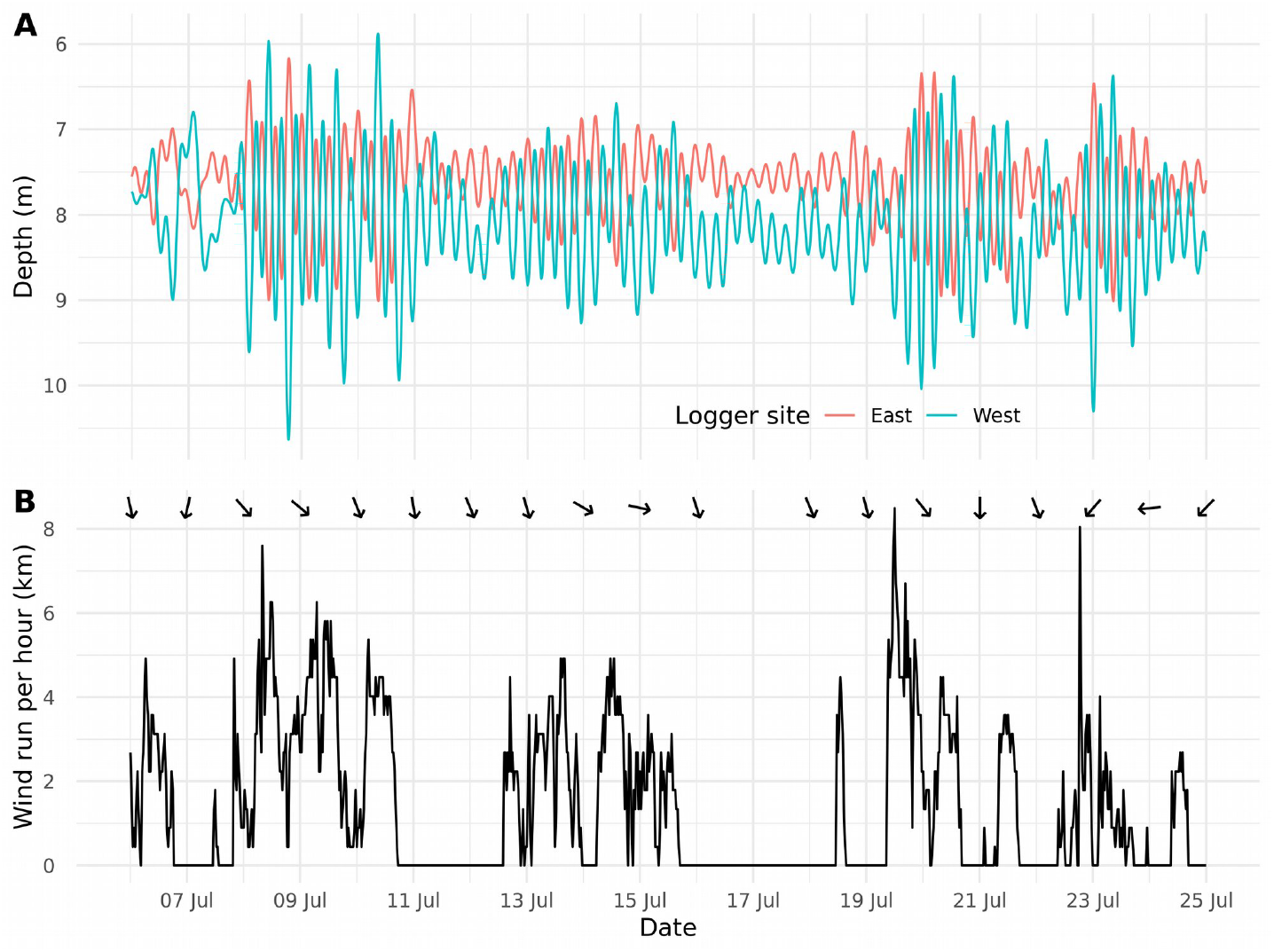
An overview of the final depth of the thermocline at both logger sites after FFT smoothing (A) and wind run (B) with mean daily wind direction (arrows) during July 2015.

### 2.1. Statistical analysis

We used Generalized additive models adopting a mixed-effects structure (GAMM) to investigate whether the amplitude of the thermocline representing strength of the seiche influences the vertical movement of fish. To assess both intra- and inter-individual variability in fish behavior we fitted a series of models including a random-effects intercept for the subject-level factor fish identification number (fishID), denoted by s(fishID, bs=’re’) in R, with or without random slope effects for each thermocline predictor, denoted by (X_ji_, fishID, bs=’re’). First, we fit randomintercept-only models (herein referred as model_fishid) comprising the raw phenotypic variance in mean fish depth which may indeed reflect varying levels of repeatable inter-individual differences in behavior (Réale et al., 2007). Next, we included time and thermocline properties (seasonal depth, amplitude and mean gradient) to account for variance due to fixed-effects (model_fixed), and, finally we add random slopes-effects for the three covariates to those models (model_BRNs), so that fish depth was modelled nonlinearly across the range of each covariate and over time. The latter implements a framework of behavioral reaction norms (BRNs) (Dingemanse et al., 2010; Dingemanse & Dochtermann, 2013) that allowed us to evaluate the behavioural responses of fish to different intensities of seiche after controlling for individual differences due to the mean temperature gradient and seasonal depth (Equation 5.2 in supporting information). A temporal reaction norm was ruled out, as the variance for the temporal fish-specific effects was negligible (σ_t,slope_ ~ 0) in all cases. We explored within-context repeatability in fish depth use across time. For each of the fitted models (species-diel period) we calculated the repeatability (*R*) index as the proportion of the total phenotypic variance (V_total_) that can be attributed to differences among fish (V_fishID_) (Nakagawa and Schielzeth, 2010). In GAMMs the smoothing components are penalized for wiggliness by including a penalty term, *β^T^S_i_β*, which is a function of the smoothing parameter (*λ*) of the model (Wood, 2008, 2017). This link can be exploited to convert the penalized regression terms into conventional random-effects (herein indicated with the argument bs=’re’) where the variance of random effects is the result of dividing the scale parameter (ϕ) by the smoothing parameter of the fitted model. Since in GAMM ϕ equals the residual variance component (*V_e_*), which can be interpreted as the intra-individual variation, and the random smoothing intercept represents the inter-individual variance (V_fishID_ = τ_00_), repeatability can be estimated as:

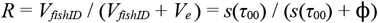

Unadjusted within-context repeatability (*R_0_*) was calculated for intercept-only models and adjusted repeatability (*R*) after controlling for confounding effects of covariates in models with fixed-effects, with random slopes included or excluded (see supporting information Supp3 for more details on variance components and repeatability estimation in GAMMs).

All data were analysed using the random intercept and random slopes models (model_BRNs) to account for the level of behavioral plasticity in responses to seiche patterns, as well as to represent trends in the mean effect. The three thermocline predictors (seasonal_depth, amplitude and mean_gradient) were preliminarily checked for collinearity, mean-centered and scaled by dividing by 2 SD units. The analyses were run separately for day and night. Diel period was estimated with the function *crepuscule()* in the R package *maptools* (Bivand & Lewin-Koh, 2014), with daytime and night defined as periods delimited by sunrise and sunset. Each subset of data was averaged to mean 5-min values to speed up computation and keep the accuracy of data points for fish positioning detections. To handle serial correlations of the model residuals, a first-order autocorrelation structure AR(1) was incorporated into the model. This structure was found suitable for our data after identification based on Akaike Information Criterion (AIC) screening of different GAM residual modelling structures and by inspection of the ACF and PACF residuals from models without autocorrelation. We used the *start_value_ rho()* function in the *itsadug* package (van Rij et al., 2017) to determine the starting lag values (Box et al., 1994). Models were fit using restricted maximum likelihood (REML) estimation with Gaussian or Gamma error distributions and identity or log link functions. Each covariate was fitted using a single penalized smoothing function based on thin plate regression splines (specified by bs=’ts’) representing an average fixed-effect across all fishID levels. GAMMs were implemented in R software version 3.6.3 (R Core Team, 2020) using the *bam()* function of the R package *mgcv* (Mixed GAM Computation Vehicle with Automatic Smoothness Estimation, *v*. 1.8-33) (Wood, 2020) (see supporting information for details of GAMM specification in R).

## 3. Results

The period under study was characterized by suitable wind frequency and strength, with the predominance of the southeast wind direction, close to the longest axis of the lake, which consequently resulted in high frequency and amplitude of internal seiche events (Figs 1–2). The partial effects of the smooth functions of the internal seiche amplitude are illustrated in Figs 3–4.

**Figure 3.**
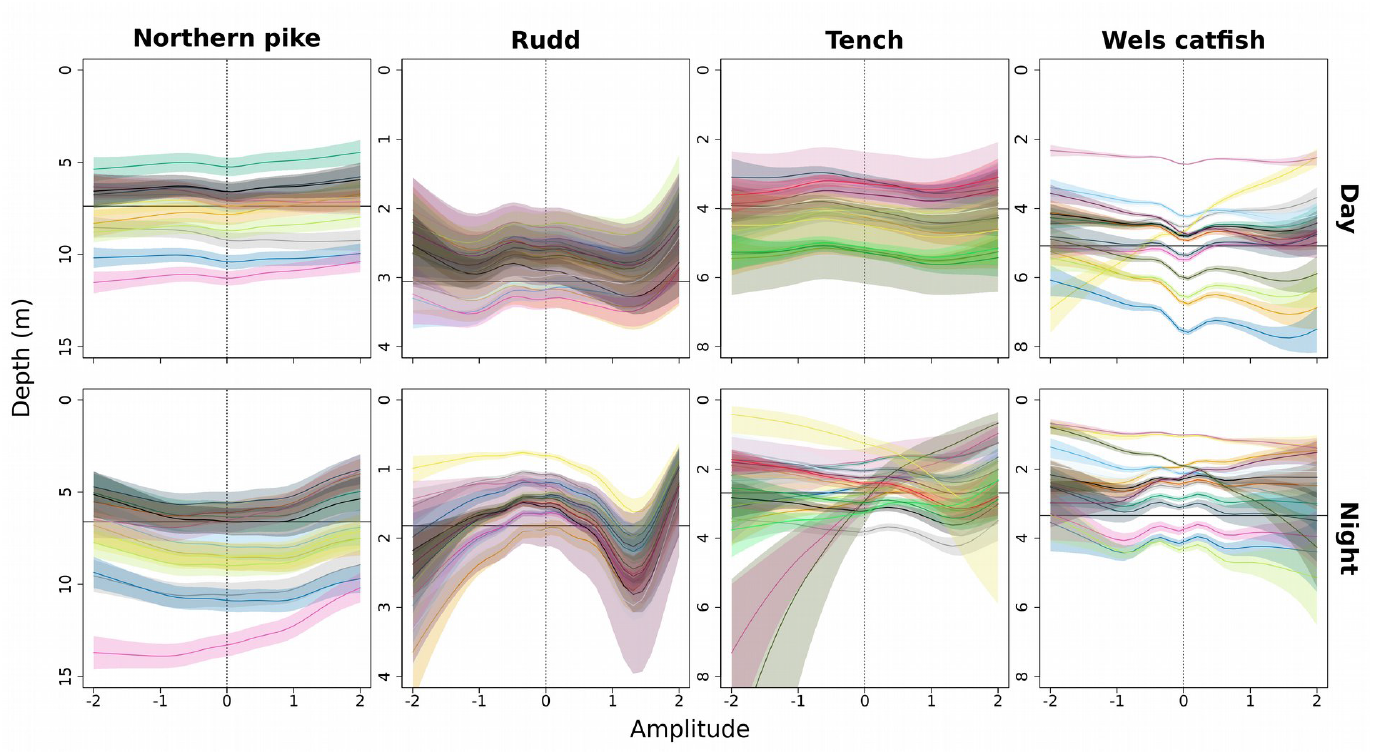
Plots of fish response to internal seiche as a function of gradual thermocline amplitude (i.e., difference between seasonal and actual thermocline depth). Depth was measured between June 16 and October 15 in 12 adult northern pike ( Esox lucius), 15 rudd (*Scardinius erythropthalmus*), 19 tench (*Tinca tinca*) and 15 wels catfish (*Silurus glanis*) during daytime and at night, with measurements averaged per 5-min intervals. Each line corresponds to an individual trajectory across increasing seiche amplitude (left to right) with zero representing periods without seiche (dotted line). Mean depth is denoted by a horizontal line. Non-linear trends were modelled as reaction norms, indicative of behavioral plasticity, by including random intercept and random slope effects in the model. The shaded area represents the 95% CI for the fitted curve, indicating the range of action of an individual or variability in a reaction norm.

**Figure 4.**
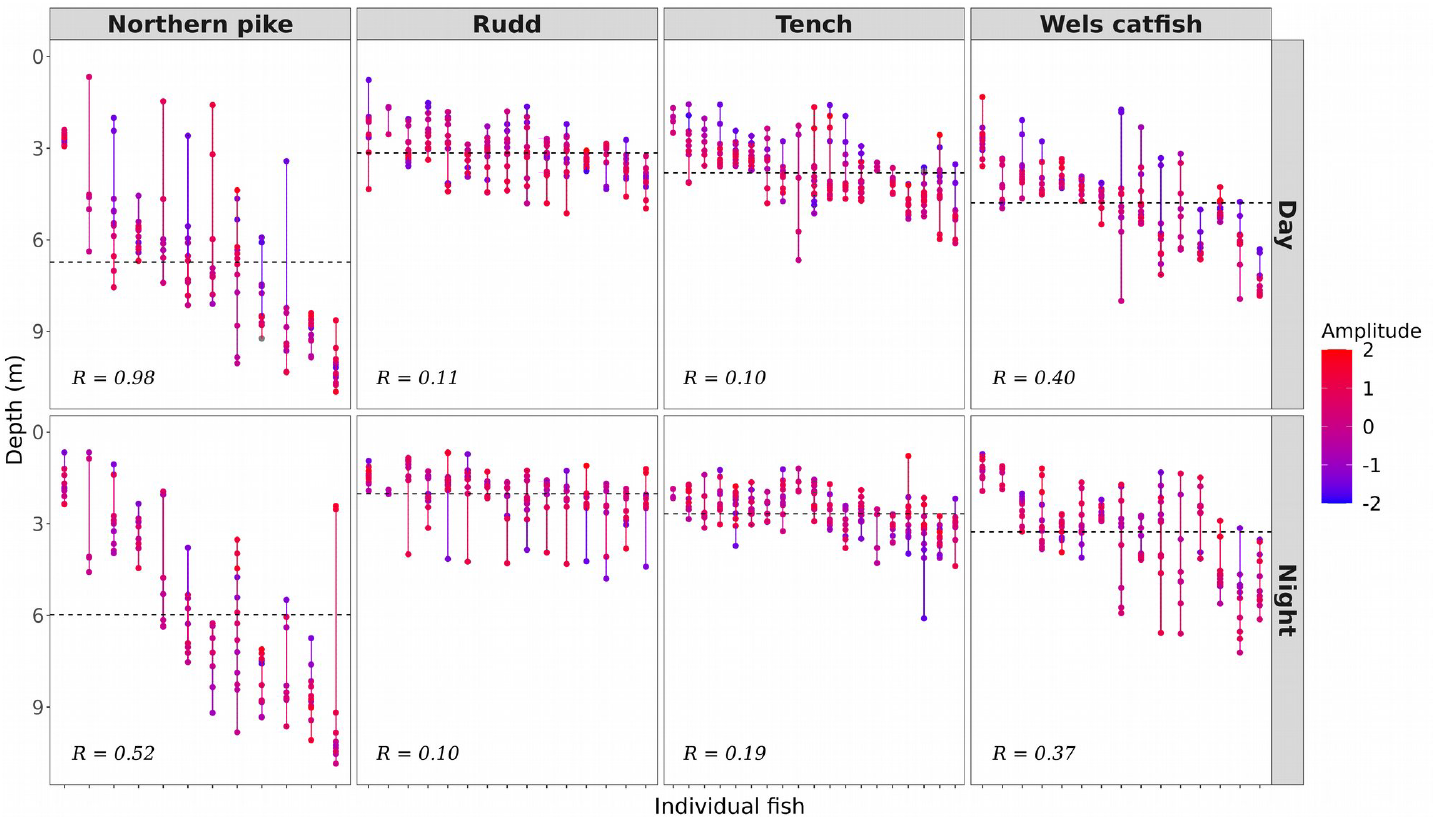
Inter- and intra-individual variation in depth use across the amplitude of seiche (depicted by color gradient) according to diel periods. Individual fish are arranged in increasing order of mean depth performance (x-axis, left to right). The average depth use of the group is denoted by a dashed horizontal line. A single vertical line denotes variation in a fish mean depth across a range of average amplitudes (i.e., individual repeatability) with a longer line indicating higher intraindividual variation. Within a group behavioral differences among fish are shown from left to right, with less overlapping vertical lines indicating overall larger repeatable behavioral differences between the individual fish. Adjusted repeatability (R) was computed from fitted GAMMs after controlling for confounding effects (fixed and random factors) of seasonal depth and mean gradient over time.

There were strong interspecific, intraspecific and diurnal differences in the patterns of fish reaction to internal seiche revealed by the model (Table 2, Figs 3–5). Even though rudd were mainly in shallow water, further away from the thermocline, the effect of seiche was significant during both day and night (amplitude_day_, F = 14.5, *p* = 0.007; amplitude_night_, F = 577.9, *p* < 0.001). Tench was mostly located just above the thermocline during the day, and reacted to upward seiche by moving to shallower water (amplitude_day_, F = 38.9,*p* < 0.001), while during the night it resided close to the water surface, and showed no reaction to seiche (amplitude_night_, F = 357.8,*p* > 0.1). The two predator species were characterized by strong individual variability. Wels catfish individuals mostly remained above the thermocline, but covered the entire depth range between the surface and the thermocline. Their movement patterns during seiche events varied among individuals, with significant reaction to seiche observed only during the day (amplitude_day_, F = 8459, *p* < 0.001; amplitude_night_, F = 217..8, *p* > 0.1). Individual northern pike showed two distinct behavioural patterns, with some individuals remaining at greater depths, well below the thermocline, while other individuals remained in the vicinity of or just above the thermocline, and showed significant reaction to seiche only during the day (amplitude_day_, F = 123.4, *p* = 0.019; amplitude_night_, F = 70.7, *p* > 0.1).

**Table 2.** Model results on the reaction of fish to internal seiche in the Lake Milada. Generalized additive mixed models (GAMM) analyzing trends in depth movement of fish according to physical properties of the thermocline (seasonal depth, amplitude and mean gradient) and over time. Numbers refer to estimates (parametric coefficients), estimated effective degrees of freedom (EDF) for main-effects smoothers, and variance components (± 95% confidence intervals, CI) for random-effects smoothers where fishID specifies random factor intercepts and random slopes for each predictor variable. Interindividual variability is modelled with similar functional responses among fish through context-dependent covariates. In all models, *s()* are smoothing functions of the covariates computed using penalized regression splines. The maximum number of knots (k-value) used to fit each model was set to 10 for all splines. The EDF indicates the degree of complexity (i.e. wiggliness) of the nonlinear relationship between a covariate and the dependent variable, depth. *Scale parameter* (ϕ), the estimated scale parameter used to calculate the variance of the i.i.d. Gaussian random effects of the penalized coefficients. *Adjusted-R^2^*, measures the goodness-of-fit of the model as the proportion of variance explained. *Deviance explained*, refers to the proportion of null deviance explained by the model. *P*-values: ^*^ *p*<0.05, ^**^ *p*<0.01, ^***^ *p*<0.001.

| Predictors | Northern Pike |  | Wels catfish |  | Tench |  | Rudd |  |
| --- | --- | --- | --- | --- | --- | --- | --- | --- |
|  | Day | Night | Day | Night | Day | Night | Day | Night |
|  | (n=12) | (n=12) | (n=15) | (n=15) | (n=19) | (n=19) | (n=15) | (n=15) |
| **Parametric coefficients** |  |  |  |  |  |  |  |  |
| (Intercept) | 6.80 | 6.25 *** | 1.56 *** | 1.06 *** | 4.08 *** | 0.95 *** | 2.99 *** | 0.52 *** |
| **Smooth terms** |  |  |  |  |  |  |  |  |
| s(seasonal depth) | 7.88 | 8.34 | 8.94 *** | 8.78 *** | 8.54 *** | 8.86 *** | 8.52 *** | 8.92 *** |
| s(amplitude) | 7.33 * | 7.53 | 8.67 ** | 8.30 | 5.86 *** | 7.38 | 7.92 ** | 8.70 *** |
| s(mean gradient) | 4.07 | 4.80 | 8.51 ** | 7.46 | 6.93 *** | 8.28 * | 7.24 | 8.13 *** |
| s (time) | 3.63 *** | 8.36 *** | 8.98 *** | 8.95 *** | 8.86 *** | 8.91 *** | 8.85 *** | 8.95 *** |
| s(fishID.seasonal depth) | 10.79 *** | 10.73 *** | 13.99 *** | 13.96 *** | 14.84 *** | 15.47 *** | 5.13 *** | 13.25 *** |
|  |  |  |  |  |  | *** |  |  |
| s(fishID.amplitude) | 9.40 *** | 9.93 *** | 13.70 *** | 13.41 *** | 9.32 ** | 16.37 *** | 9.55 *** | 12.15 *** |
|  |  |  |  |  |  |  |  | ** |
| s(fishID.mean gradient) | 8.66 *** | 9.53 *** | 13.41 *** | 13.04 *** | 11.90 *** | 16.63 *** | 9.96 *** | 12.78 *** |
| s(fishID) | 10.79 *** | 10.81 *** | 13.99 *** | 13.99 *** | 15.99 *** | 16.64 *** | 12.59 *** | 13.32 *** |
|  |  |  |  |  |  | *** |  |  |
| **Scale parameter ($\Phi$)** | 3.86 | 4.63 | 0.09 | 0.32 | 2.14 | 0.27 | 0.96 | 0.69 |
| **Adjusted R<sup>2</sup>** | 0.54 | 0.56 | 0.59 | 0.56 | 0.39 | 0.19 | 0.35 | 0.33 |
| **Deviance explained (%)** | 53.8 | 56.5 | 50.4 | 48.3 | 38.7 | 19.1 | 35 | 33 |

**Figure 5.**
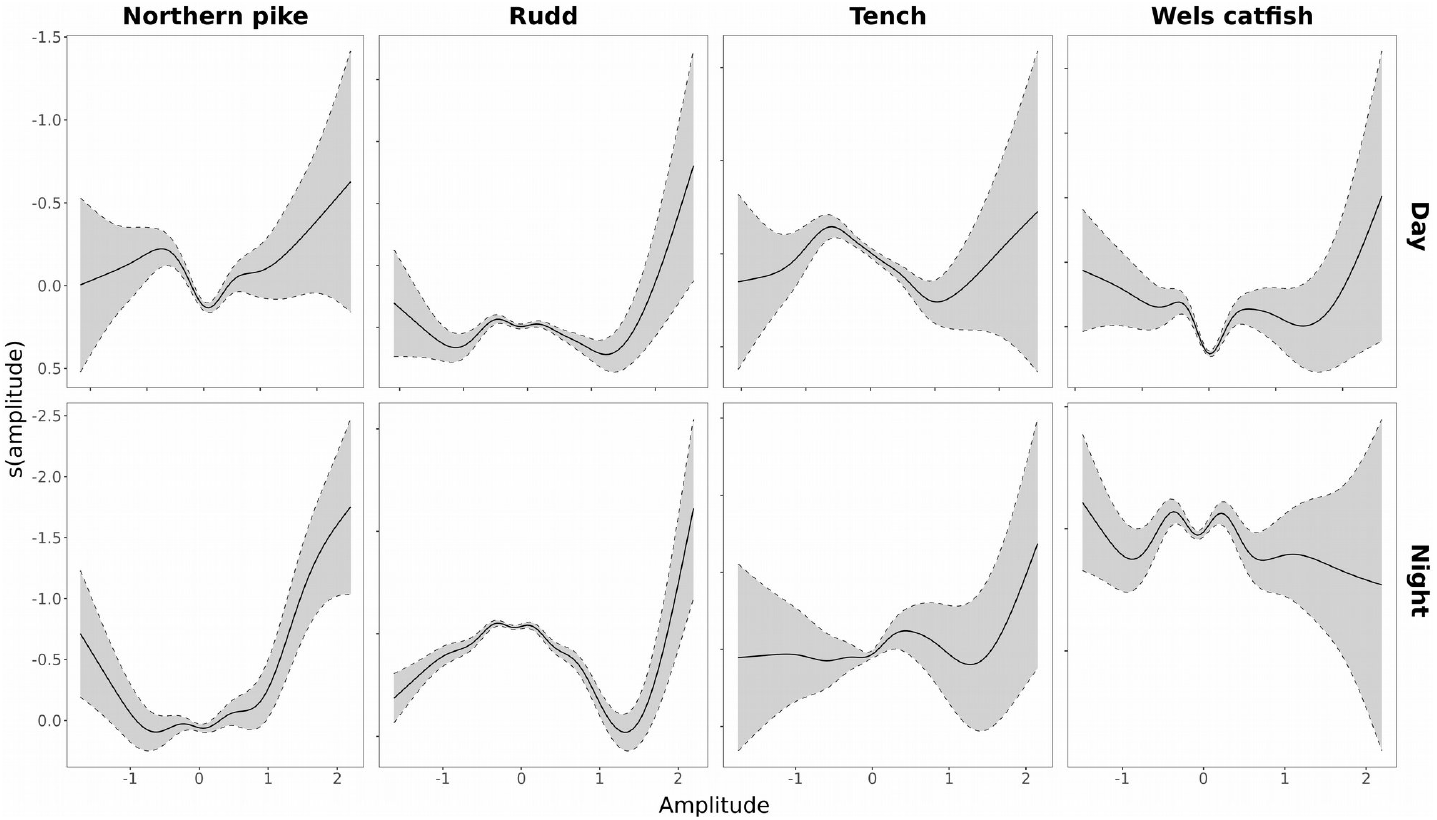
Partial effects of the smooth functions of the amplitude of seiche on fish vertical movement of northern pike, rudd, tench and wels catfish according to diel periods. The slopes represent average effects for the amplitude of thermocline based on GAMs fit to 5-min average fish position data over a whole time period. Splines for other predictors are omitted.

The final BRNs models, variance components and the effects of smooth functions of each covariate on depth are shown in Table 2. The performance of the models was highest for northern pike during the day (Adj R^2^ =0.59) but remained relatively low for all models, which is reasonable given the high variability over the whole time series. Effective degrees of freedom (EDF) showed consistency between diel periods in all models, except for tench, where smooth amplitude functions were less wiggly during the day than at night, indicating less variability both in terms of the mean seiche effect and the level of plasticity and response to seiche patterns. On the other hand, the mean_gradient smooth functions in northern pike showed relatively low variability compared to the other species.

In our behavioral reaction norm approach we accounted for the relationship between thermocline properties and repeatable inter-individual differences in average behavior across time (i.e., personality), and individual responses to each context (i.e. plasticity), especially those driven by seiche intensity. Reaction norms are subject to different seiche intensities (Fig. 3) and reflect well the level of repeatability and depth variation of fish (Fig. 4) during daytime and nighttime activity. In fact, depth was significantly repeatable across all four species, as shown by the nonoverlapping 95% confidence intervals for the variance components of the random effects (Tables 2 and 3). In particular, repeatability across thermocline gradients was low for tench and rudd during both diel periods (tench *R_day_*= 0.19, R_night_= 0.19; rudd tench *R_day_*= 0.11, R_night_= 0.10), moderate for wels catfish (Rday= 0.40, Rnight= 0.37) and very high for northern pike during the day (*R_day_*= 0.98) and moderate at night (*R*_night_= 0.52).

**Table 3.**
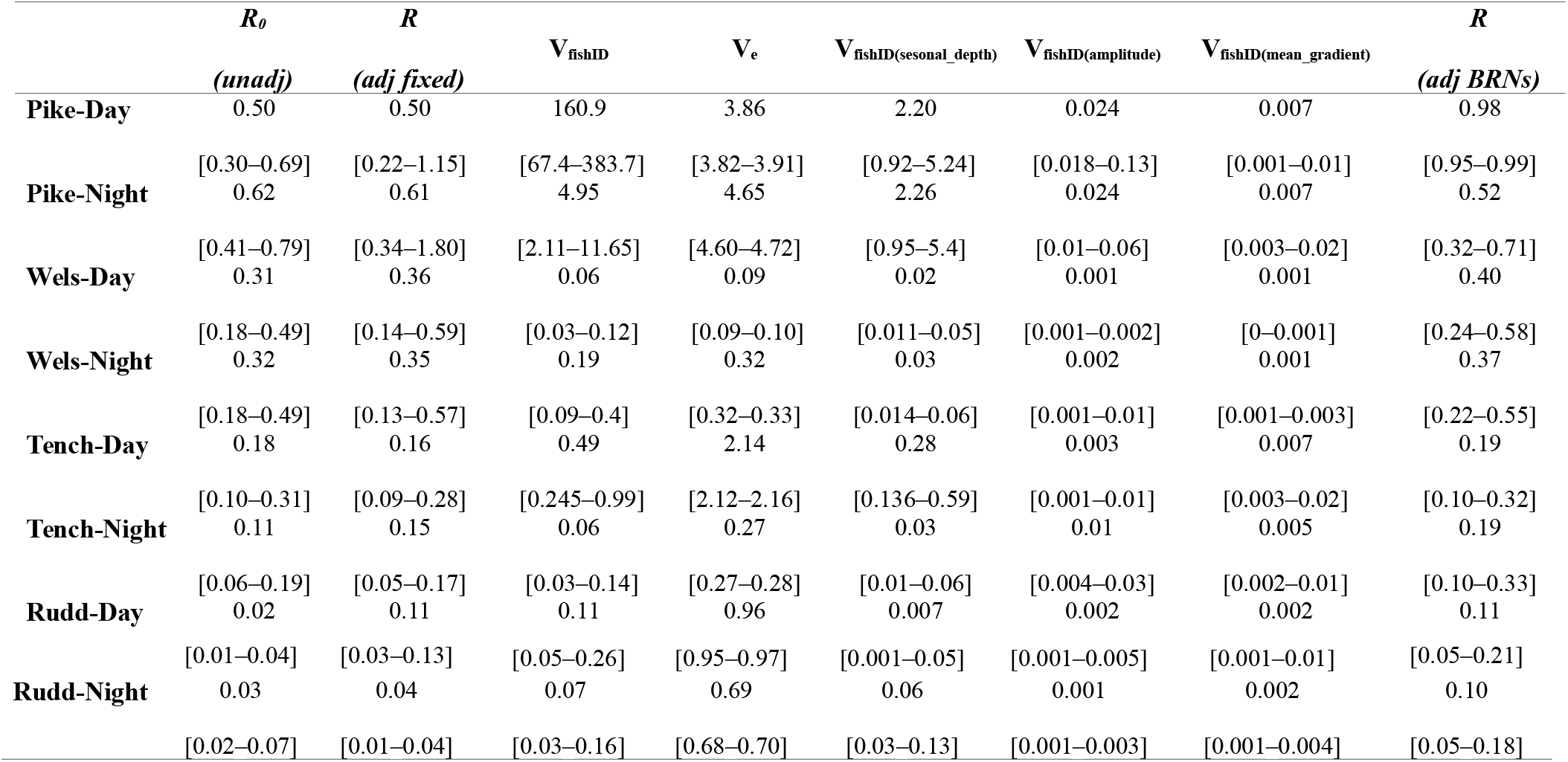
Repeatability in individual behavior across contexts of thermocline –dependent variables. Data show adjusted and unadjusted *R* values with 95% confidence intervals (CI) calculated from model_fishid, model_fixed, and model_BRNs GAMMs. Variance components are also shown for model_BRNs only. Random effect variances for each covariate are calculated by dividing the estimated scale parameter (ϕ) by the smoothness parameter (λ) of the fitted GAMM. *R*epeatability was calculated using a nonlinear approximation, according to the definition of Nakagawa and Schielzeth (2010) as the ratio of the interindividual variance (V_fishID_), given by the random smoothing intercept s(fishID) (i.e., *τ*_00_), and the total phenotypic variance (V_total_ = V_fishID_ + V_e_), where V_e_ is the residual variance of random effects (or intraindividual variability) and is equal to ϕ (see SM for more details on variance components and estimation of *R*).

When compared to simpler models (model_fishid, model_fixed), these results showed that the inclusion of slopes for each thermocline covariate primarily affects repeatability estimates for northern pike, as *R* increases considerably by almost 50% during the day and by 10% during the night. However, there are important differences between the two data sets. During the day, interindividual variation in behavior is much greater than during the night (τ00 ~ 160.9 versus τ00 ~ 4.95), indicating a greater influence of thermocline environmental variables, including the reaction of fish to seiche, during the day. For other species, these environmental variables, regardless of their effects on individual behavior, did not cause significant changes in behavioral variability among individuals, except for tench during nocturnal activity (Table 3).

## 4. Discussion

Our study showed that internal seiche affects fish behavior. Except for rudd, fish reacted to seiche only during the day, and the reaction was mostly absent at the smallest seiche amplitudes, manifesting itself only during the strongest seiche events (Figs 3, 5). Reactions were also considerably stronger in the presence of upward than downward seiche events. This was most likely because the upward movement of the thermocline led to a narrower vertical depth range of the epilimnion, thereby compressing the epilimnetic habitat which the studied species normally inhabit. Conversely, the downward movement of the thermocline merely provided a transient expansion of the epilimnion depth range. Individual variation in reaction to seiche dynamics and depth preference was stronger for predatory species, northern pike and wels catfish, than for generalist species, rudd and tench. The effect of seiche was weak compared to the impact of thermocline, but was nevertheless significant.

The study represents the first attempt to address this question at such a high level of spatiotemporal resolution, provided by fine-resolution fish telemetry. The rare examples of studies with a similar scope, focusing on the effects of internal seiche on the vertical distribution of lacustrine fish, were typically based on hydroacoustic transects (Levy et al., 1991; Easton & Gophen, 2003; Bass et al., 2014). For example, Levy et al. (1991) detected altered rates of vertical positions of juvenile sockeye salmon (*Oncorhynchus nerka*) during internal lacustrine seiche events, whereas Easton and Gophen (2003) found only a weak effect of seiche on fish distribution. Other studies on internal seiche dynamics focused only on hydrological aspects (e.g. Cossu et al., 2017) or on the resulting fish population dynamics, such as effects on fish egg survival and resulting year-class strength (Aalto & Newsome, 1993).

The impact of internal seiche dynamics on lacustrine organisms depends not only on temperature, but also on oxygen variability, as anoxic conditions may occur below strong thermoclines (Vejřík et al., 2016; Cossu et al., 2017). However, as hypoxic conditions did not manifest in the lake within the range of seiche influence during the present study (Říha et al. 2021), the observed effects of seiche were likely manifested only through the temperature stress. Temperature variability is considered a key factor and stimulus affecting fish movement (Capra et al., 2018), and internal seiche events are capable of rapidly affecting the vertical distribution of temperature (Levy et al., 1991). In addition to direct effects, internal seiche dynamics can also produce indirect effects on fish distribution by affecting the distribution of their prey (Easton & Gophen, 2003; Aspillaga et al., 2017). However, we were unable to discriminate such effects in the present study, as reaction of zooplankton, mobile benthic invertebrates (i.e., prey of rudd and tench, Vejříková et al. 2016) or whole fish prey community (prey of northern pike and wels catfish) were not sampled during the study.

The reaction to internal seiche dynamics differed among species. However, it only partly follows our hypothesis. The weakest reaction was as expected found for northern pike, but the other species had the same reaction during the daytime and only rudd (temperature-tolerant species) reacted during the night time. Observed inter-specific differences can be linked to species-specific ecology, diurnal activity patterns, trophic level, and foraging strategies, but especially to their thermal preferences. Optimal temperature and thermal tolerance ranges are species-specific and can vary widely, including thresholds that can lead to cold-shock stress and various physiological and behavioural responses (Donaldson et al., 2008).

The effects of seiche on northern pike were expected to be weak, considering that it is a cool-water predator, flexible in its responses to varying habitat conditions (Craig, 2008). The ability to cope with ambiguous environmental temperatures was also demonstrated in our study, where individual depth preference varied widely, with individuals distributed in both deeper (majority of tracked fish) and shallower waters. Despite this, even northern pike reacted to seiche-induced movement of the thermocline and adjusted depth accordingly (i.e., individuals preferred deeper habitats, Fig. 4) during the day. Individuals located in deeper water preferred to remain in the vicinity of the thermocline and their response to seiche could be more a maintenance of position in the thermocline than a reaction to temperature changes (as the northern pike’s reaction pattern was the closest to linear, reacting even to downward shift). On the other hand, it is evident that northern pike are less affected by low temperatures than the other three species studied and is therefore less susceptible to cold shock from a shifting thermocline. Further, northern pike is a diurnal species with low nocturnal activity (Baktoft et al., 2012). It can also explain weaker patterns in reaction to seiche during night.

Wels catfish prefers warmer water temperature (Capra et al., 2018; Cucherousset et al., 2018), and as such their reaction to seiche events by avoiding cooler hypolimnetic water was expected. However, the model did not detect such an effect at night when wels catfish showed no response to seiches events. Diurnal differences can be associated with differences in preferred depth between day and night, or with the diel activity patterns of the species. Wels catfish occupied lower depths during night, and as such they were less exposed to seiches dynamics. Although wels catfish is a nocturnal species (Pohlmann et al., 2001; Vejřík et al., 2017), its activity can vary with the seasons, and during warmer periods it tends to also be active during the day (Slavík et al., 2007; Cucherousset et al., 2018). Wels catfish could also change their movement strategy between day and night as they are characterized by flexible behavioural patterns and the ability to develop a wide range of strategies (Vejřík et al., 2017; Cucherousset et al., 2018).

Studies on the ecology of tench and rudd have been relatively scarce and much is still unknown about their habitat preferences, feeding ecology and behaviour. Tench is considered an eurythermal species (Heap et al., 1985), with a high level of tolerance to environmental conditions such as low oxygen concentrations, relatively high salinity, and high water temperature (Weatherley, 1959; Kottelat & Freyhof, 2007). However, it exhibited a significant reaction to seiche dynamics. According to Rosa (1958 cit. in Weatherley, 1959), tench avoids extreme cold temperatures, so it is possible that, although highly adaptable to different water temperatures, it is still susceptible to rapid temperature changes and cold shock. Interestingly, it is considered a strictly nocturnal species (e.g. Herrero et al., 2003), whereas in the present study it was shown to react to seiche dynamics with vertical movements only during the day.

Rudd is a mainly littoral species, herbivorous or planktivorous depending on size and temperature, and tolerant to a wide range of temperatures (García-Berthou & Moreno-Amich, 2000; Hicks, 2003; Kottelat & Freyhof, 2007; Vejříková et al., 2016). In the present study, it was affected by both daytime and nighttime seiche dynamics, which is consistent with previous reports of its bimodal activity, manifested around noon and midnight (Hohausová et al., 2003). This reaction was probably driven by the strongly temperature-dependent physiology of plant matter digestion by rudd, the major part of the rudd diet in the lake (Vejříková et al., 2016). Rudd digest plant matter with the help of intestinal microflora capable of utilizing cellulose only in relatively warm water, above 17 °C, which could have prompted rudd to react to seiche to avoid exposure to cold water (Vejříková et al., 2016).

The predatory species in this study, northern pike and wels catfish, showed significant and repeatable differences in the use of depth in the water column context, while these differences were not as consistently repeatable for generalist species, rudd and tench. As a result, inter-individual differences in behavior were greater for predatory species while they were reduced in generalist species. We see therefore that, regardless of consistent differences in individual behavior within each of the four species, on average, predators tended to change their depth occupancy more frequently in response to internal seiche than generalists (Figs 4–5).

Statistical models used to test the effect of seiche dynamics on fish depth include seasonal depth, mean gradient and amplitude. However, the effect of amplitude, representing seiche strength, was difficult to isolate, given that it is strongly interrelated and dependent on the other two parameters. Therefore, our investigation of inter-individual variation in depth-use behaviour is limited to a single open context, that of the water column, rather than to the fringe where the internal seiche typically occurs. These differences might be associated with prey distribution. Tench and rudd forage mainly on benthic prey or palatable macrophytes which are more abundant in shallow littoral (Vejříková et al., 2016) while northern pike and wels catfish have plastic foraging strategies including larger number of prey types (Vejřík et al., 2017). In addition to that, the risk of predation could also play an important role. Tracked individuals of northern pike and wels catfish were adults at low risk of predation by conspecifics or other predators and, therefore, were able to use a wider range of depth habitats. Studied rudd and tench individuals were also adults, but they were still under the potential risk of predation by northern pike and wels catfish (Vejřík et al., 2017), while beds of vertically well-developed macrophyte species that serve as shelter (as *Myriophyllum spicatum* or *Potamogeton pectinatus*) were only present up to 4-6 m of depth (Vejříková et al., 2021). However, more detailed research is needed to understand interaction between individual traits (morphological, behavioural, or physiological) and seiche dynamics.

It is important to note certain limitations of the study. The approach to inferring the presence of internal seiche was relatively simple, as it was based on data from only two logger sites in the lake, which did not allow for more precise tracking of seiche dynamics at higher resolution. Our study was also based on a relatively small number of fish tracked (Table 1), and did not investigate juvenile stages or the possible effects of ontogeny on the reaction of fish to seiche, nor did it consider the effects of fish sex or physiological state. Furthermore, as only a minor part of the lake populations were studied, we were unable to take into account social aspects and the effects of inter- and intra-specific interactions. Fish telemetry studies face a number of common challenges, which should be taken into account both in the planning phase of the project and in the interpretation of the results (Brownscombe et al., 2019; Lennox et al., 2021).

The strength of the internal seiche depends mainly on wind speed and duration, but in elongated lakes the wind direction may in fact be the key factor. For example, in the Lake Rehtijarvi (Finland), the greatest seiche amplitude was manifested when the wind direction coincided with the longitudinal axis of the lake, even if the wind was not particularly strong during those periods (Horppila & Niemistö, 2008). Other relevant factors that can affect seiche dynamics are the shape, size, and depth of the lake, and the topography of the lake bottom and surrounding terrain (Aalto & Newsome, 1993).

Internal seiche dynamics is able to affect aquatic communities. Seiche events have recognized ecological effects (Wells & Parker, 2010), with a potential to affect fish community structure, dynamics, and stability in several ways. First, they may affect the survival of eggs, embryos, and juvenile fish, and cause spatial variation in survival and recruitment (Aalto & Newsome, 1993). They could also play an important role in egg depth deposition and egg incubation time (Čech et al., 2012). Internal seiche has also been found to weaken the association of fish larvae and zooplankton with the thermocline in marine environments compared to periods without seiche, which could affect biotic interactions (Gray & Kingsford, 2003). Second, they may affect physiological processes and alter the behaviour of fish communities (Cossu et al., 2017). Third, thermocline shifting may induce a transient reduction in habitat for seiche-responsive species, thus potentially affecting interspecific interactions, which was already shown to be one of the effects of seiche-induced hypoxia (Kraus et al., 2015). Consequently, thermocline shifts may also affect fishery catch rates as well as the reliability of community sampling activities.

Climate change is expected to affect physical properties of lakes, including their hydrology (O’Reilly et al., 2003). It may result in warmer epilimnetic water and stronger thermoclines, and potentially affect internal seiche dynamics (Cossu et al., 2017). How these changes in internal seiche will manifest will depend primarily on particular regional and local changes in wind patterns and intensity (Pryor et al., 2005; Tobin et al., 2015). For example, reduced wind speed in the Lake Tanganyika area, attributed to climate warming, has contributed to reduced water mixing and upwelling (O’Reilly et al., 2003). In addition to trends in mean wind speed, seiche dynamics are likely to be particularly affected by the frequency of extreme winds and storms, which are expected to intensify with climate change (Seidl et al., 2017). Furthermore, climate change may also indirectly reinforce the impact of seiche events through water warming, as warmer surface layers are likely to increase the susceptibility of fish to cold shock from exposure to hypolimnetic water (Cossu et al., 2017).

In conclusion, our study demonstrates that short-term thermocline dynamics have an effect on the distribution of fish communities. Although the consequences of internal seiche on lacustrine fish communities have been poorly studied and understood, they are especially likely to gain in importance with ongoing climate change (Donaldson et al., 2008). Further research is needed to investigate the relationship between physical and biological factors related to internal seiche dynamics and, especially, for studies to compare their importance across multiple stratified lakes with a wide range of conditions (Cossu et al., 2017). Further studies should also focus on the possible effects of internal seiche dynamics on fish distribution and horizontal movement.

## Supporting information

S1 - Information on tag type and tagging procedure

S2 - Filtering Umap Positions

S3 - Extended description of the model

## Supporting information

Additional supporting information may be found online in the Supporting Information section at the end of the article.

## Data Availability Statement

The analyzed dataset is available on GitHub:

https://github.com/allantsouza/ms-jaric-2020-fish-reaction-to-seiche/tree/develop/data/detections

The scripts used for analysis are available on GitHub:

https://github.com/allantsouza/ms-jaric-2020-fish-reaction-to-seiche/blob/develop/outputs/seiche_ms_v5.md#headamodel_BRNs

## Acknowledgements

IJ acknowledges the sponsorship provided by the J. E. Purkyně Fellowship of the Czech Academy of Sciences, as well as support by the Alexander von Humboldt Foundation. Authors acknowledge funding provided by the European Union’s Horizon 2020 research and innovation programme through the project ClimeFish (grant No. 677039). The work was also supported from ERDF/ESF project Biomanipulation as a tool for improving water quality of dam reservoirs (No. CZ.02.1.01/0.0/0.0/16_025/0007417) and the project QK1920011 “Methodology of predatory fish quantification in drinking-water reservoirs to optimize the management of aquatic ecosystems”. The authors thank Finn Økland for support during field work, and Marie Prchalová and Lobsang Tsering for helpful comments on an earlier version of the manuscript.

## Authors’ contributions

IJ, MR, and VD conceived the ideas and designed methodology; JP, MR, KØG, PB, MH, TJ, MM, ZS, MŠ, LV, IV and VD conducted the acoustic telemetry study and collected the tracking data. VD, MR and KØG processed the data. ATS and RRB performed statistical analysis. IJ led the writing of the manuscript. All authors contributed critically to the interpretation of results and manuscript drafts and gave final approval for publication.

