## Supplementary material for "Influence of internal seiche dynamics on vertical movement of fish": S1 - Information on tag type and tagging procedure

**Electronic Supplementary Material S1**

### Information on tag type, dimensions and weight, and tagging procedure (anesthesia and surgery) per species

| Species | Tag type | Tag dimensions (mm) | Tag weight (g) | Anesthesia conc. ($ml*L^{-1}$) | Time in anesthesia (min) | Surgery duration (min) |
|---|---|---|---|---|---|---|
| Pike | MM-M-11-28-PM | 65x12 | 13 | 0.7 | 3.6 | 2.8 |
| Rudd | MM-M-8-SO-PM | 47x8.5 | 6 | 0.7 | 1.75 | 3.95 |
| Tench | MM-M-11-SO-PM | 47x11 | 8.2 | 0.8 | 2.3 | 3 |
| Wels | MM-M-11-28-PM | 65x12 | 13 | 1 | 4.5 | 3 |
