## Supplementary material for "Influence of internal seiche dynamics on vertical movement of fish": S2 - Filtering Umap Positions

**Electronic Supplementary Material S2**

### Filtering of positions estimated by the U-MAP software

U-MAP (User Managed Acoustic Positioning) is a proprietary software from Lotek Inc. that processes transmitter detection data collected with autonomous Lotek WHS-receivers. The detection data recorded by each WHS receiver include tag ID, time of arrival (TOA) and relative signal strength or power of the detected signal. U-MAP uses these data to estimate transmitter positions by means of multilateration on TOA data for transmissions that were detected at three or more receivers. The algorithms solve non-linear equations involving hydrophone positions and the difference in time of signal arrival (TDOA) at each hydrophone detecting the transmission. Depending on the number and geometry of the hydrophones involved in the position calculation, the equation may have multiple solutions (Lotek 2014). These are referred to as twin solutions (or shadow solutions). U-MAP saves all twin solutions to the output file, and it is up to the user to choose, or filter, the correct position between the twin positions. Moreover, the use of TDOA implies hyperbolic equations, which also tend to give large errors for certain receiver and transmitter configurations. These configurations depend on the receiver array and the transmitter location, but also on stochastic variation in which receivers detect the signal. Two successive position estimates may therefore differ in location even when the transmitter did not move, if the receivers involved in the position estimate differ. Position estimates produced by programs using TDOA-methods, such as U-MAP, therefore require filtering by post-processing in order to eliminate erroneous position estimates.

The performance of the position estimates and potential filters were evaluated by short-term tag-tows from boat with high-precision GPS-device above the towed transmitters, by several stationary reference tags throughout the study (Fig. S1), and by visual inspections of all fish tracks and filtered results (Fig. S2).

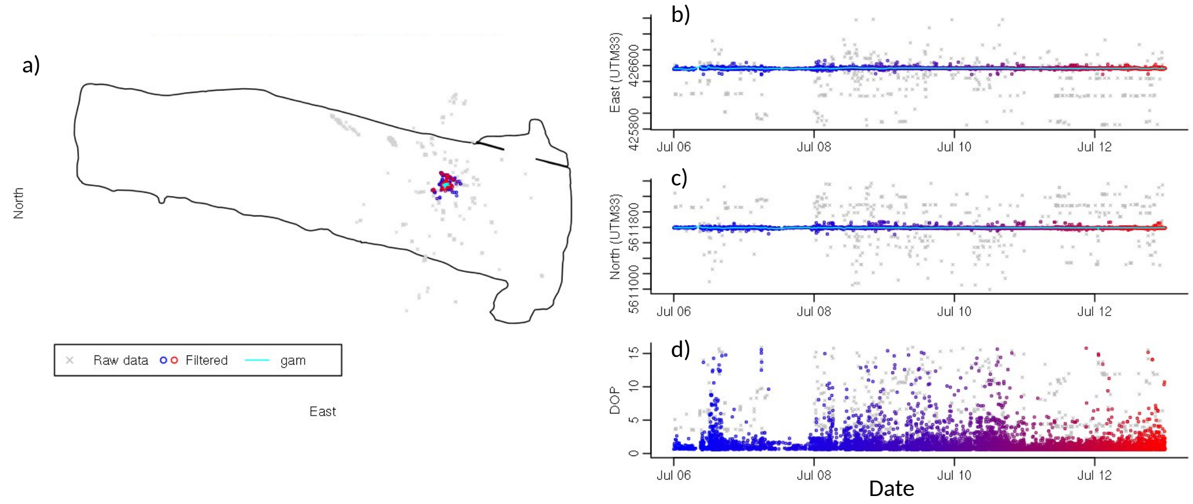


*Figure S1. Example of filtering one week of position data for a reference tag. Grey colour indicates positions not accepted by the filtering algorithm, blue to red indicates accepted positions with colour indicating time; blue being oldest and red being latest positions. The cyan-coloured line indicates the final gam-smoothed track. a) Horizontal position estimates, with the outline of Lake Milada indicated by a black line, b) easting of positions versus time, c) northing of position versus time, d) dilution of precision (DOP), a precision measure produced by U-MAP indicating the effective precision of a position estimate given hydrophone geometry and the time measurement resolution of the receiver.*

The filtering procedure required several steps to ensure the best error exclusion:

1. Calculation of detection rate $D_r$ and detection mean power $D_p$ at each receiver by 20 minute intervals for the transmitter in focus. Detection rate and detection power are expected to be highest closest to the receiver (Gjelland & Hedger, 2013). These two measures were then combined into one receiver signal scale $D_s$ by scaling each of them before adding them together; $D_s = scale(D_r) + scale(D_p)$. Scaling here means centring and dividing by the standard deviation.
2. Calculation of distance weights $D_w$ for later use in calculation of regression weights, by the use of the logistic function: $D_{w,i} \frac{1}{1+2\cdot 0.001^{(D/1500-0.6)}}$, where $D_i$ is distance from the receiver $i$ to the position estimate. Parameters where subjectively fitted to obtain a desired logistic shape with high weights to receivers closer than 500 m to the position estimate, and low weight to receivers farther than 1000 m away from the position estimate.
3. Calculation of regression weights $R_w$ were then done for each U-MAP position estimate, by first ranking all receivers from highest to lowest $D_s$ value within the current 20 minute period. The $D_s$ for the highest ranked receiver was then used together with the distance weight $D_w$ for this receiver to obtain $R_w$ through the function $R_w = D_w(1 + D_s)$
4. Lake shoreline exclusion: All position estimates farther than 50 m outside the lake shoreline polygon were excluded from further filtering, and marked as false position estimates.
5. Calculation of gam-models (generalized additive model; Wood, 2011) and gam-predictions for east and north-directions. This was done for successive periods of six hours, with one extra hour of data before and after the end of each period included in the gam-regression to stabilize the ends. The resulting gam-models were then used to predict

position ($East_{gam}$, $North_{gam}$) within the six-hour period at the same time points as the U-MAP estimated positions. Gam residual was calculated as the distance between the U-MAP position estimate and the gam prediction, including residuals for positions excluded from the gam regression. The gam-formulation was y ~ s(time), where y was either east or north, and s was the smooth term specified with a cubic spline regression model. *K*, the dimension of the basis used to represent the smooth term, was set as a function of the *n* observations included in the regression period, with *K* as the nearest integer to *n*/15, but with a minimum of value of 3.

6. For U-MAP twin positions (i.e. positions with equal timestamp), the position with the lowest gam-residual was flagged as a true position, the other as false.
7. In order to remove heavy outliers with a strong influence on the gam-regression, U-MAP positions with gam-residuals > 1000 m were flagged as false.
8. A second gam-regression was now performed, repeating point 5 but excluding all positions now flagged as false. In addition, positions with gam-residuals > 150 m were excluded from the input data to the second gam regression. New gam-predictions were made for all time points of U-MAP position estimates.
9. For U-MAP twin positions (i.e. positions with equal timestamp), the position with the lowest gam-residual was flagged as a true position, the other as false.
10. Positions with gam-residual larger than 100 m were flagged as false. U-Map position estimates, gam predictions and gam residuals were stored together in the database, such that the gam residual threshold for position acceptance could later be modified if desirable.

The amount of positions varied over time, as well as the relative amount of poor position estimates. This is easy to handle for a tag with known position (Fig. S1a-c), but when the true position is not known it is necessary to rely on filtering methods using information from the detection data. The DOP measure provided by U-MAP had poor association with the position quality, as position estimates with low DOP-value could be far from the true position, and positions estimates with high DOP-value could be close to true position (Fig. S1d). The filtering procedure described above did a great job in sorting out good from bad position estimates, and in general kept > 85 % of the positions flagged as true positions.

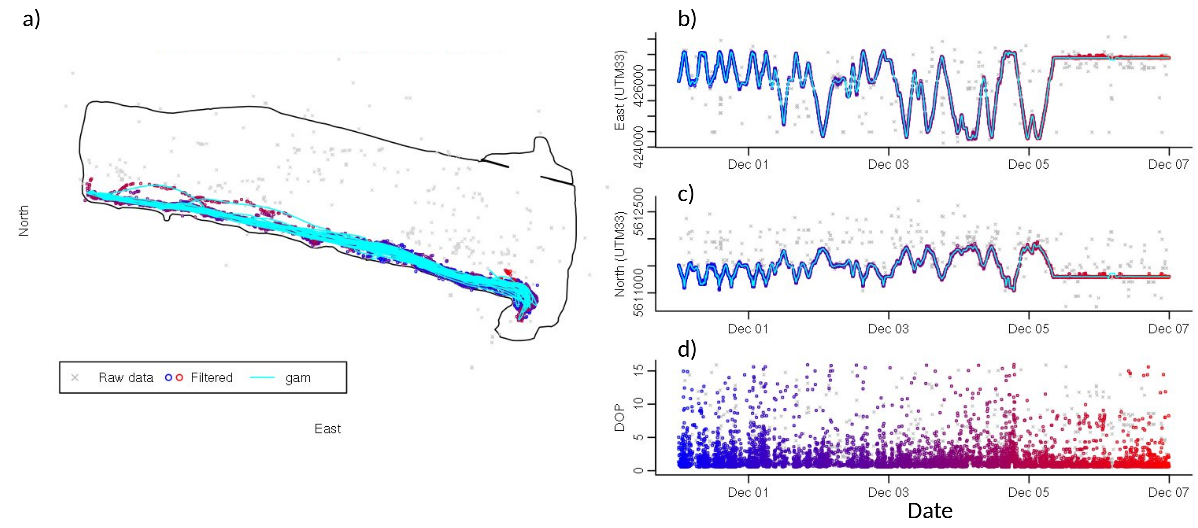


*Figure S2. Example of filtering one week of position data for an European wels individual. Grey colour indicates positions not accepted by the filtering algorithm, blue to red indicates accepted positions with colour indicating time; blue being oldest and red being latest positions. The cyan-coloured line indicates the final gam-smoothed track. a) Horizontal position estimates, with the outline of Lake Milada indicated by a black line, b) easting of positions versus time, c) northing of position versus time, d) dilution of precision (DOP), a precision measure produced by U-MAP indicating the effective precision of a position estimate given hydrophone geometry and the time measurement resolution of the receiver.*
