## Supplementary material for "Influence of internal seiche dynamics on vertical movement of fish": S3 - Extended description of the model

**Electronic Supplementary Material S3**

### Extended Methods

#### *1.1. Model definition*

Let $y_i$ be a response variable with ith observations and j covariates $x_j$ associated with fixed effect, $\theta$ a c x 1 vector of variance components and $b$ a q x 1 vector of random effects, $y_i$ is assumed to be conditionally distributed with mean $E[y_i] = \mu_i^b$ and variance $var = \upsilon(\mu_i^b)\phi$, and the random effects $b$ are assumed to be independent and identically distributed (i.i.d.) Gaussian with mean $E[b] = 0$ and variance $\Sigma_\theta = \phi\,\lambda^{-1}$, where υ is a variance function with value 1, ϕ is a scale parameter, $\Sigma_\theta$ is an identity matrix and $\lambda$ is the smoothing parameter of a penalty term $\beta^T S_i \beta$ used to penalized the log-likelihood function of the model. Assuming a semiparametric model we can write a generalized additive mixed model (GAMM) formula as follows:

$$g\left(\mu_i^b\right) = X_i\beta + \sum_j L_{ij} f_j\left(x_{ji}\right) + Z_i\left(z_i^T\right)b, b\, N\left(0, \Sigma_\theta\right), y_i\, EF\left(\mu_i^b, var\right) \quad (1)$$

, where:
$g()$ is a known link function,
$\beta$ are estimates of model coefficients,
$L_{ij}$ are linear functional terms of the linear predictors,
$X_i$ is the ith row of a p x p design matrix of fixed effects associated with covariates $x_{ji}$ that are linearly and non-linearly (by means of $f_j$) related to $y_i$,
$Z_i$ is the ith row of a p x q precision matrix of random effects b, where $z_i$ are covariates associated with random effects,
$f_j()$ are non-linear smooth functions of the covariates $x_{ji}$, represented by $A_i$ sum of $k$ simpler basis functions $b_{j,k}$ fitted to each covariate $x_{ji}$ and a vector $\gamma$ of all basis coefficients $\beta_{j,k}$:

$$f_j\left(x_{ji}\right) = \sum_k \beta_{j,k} b_{j,k}\left(x_{ji}\right) = A_i\gamma \quad (2)$$

The model is fit by maximizing a penalized log-likelihood function ($L'$) where wiggliness is penalized by adding to each smooth function a penalty term, $\beta^T S_i \beta$, dependent on a specific identity penalty matrix $S_i$ and the vector of model coefficients $\beta$. Multiplying each penalty term by a smoothing parameter $\lambda$ is a means to control the trade-off between wiggliness penalization and the model log –likelihood ($L$):

$$L' = L - \lambda \beta^T S_i \beta \quad (3)$$

Applying the Equation 1 to our own analysis and considering that the relationship between fish depth and thermocline predictors is non-linearly modelled as separate non-parametric terms (but the intercept), a general model formula can be expressed as:

$$g(E[y_i]) = \beta_0 + \sum_j f_j(x_{ji}) + Z_i b, b N(0, \phi \lambda^{-1}), y_i EF(\mu_i, \varnothing) \quad (4.1)$$

Or if model fit with a Gamma distribution and a log link function:

$$g(\log(E[y_i])) = \beta_0 + \sum_j f_j(x_{ji}) + Z_i b, b N(0, \phi \lambda^{-1}), y_i Gamma(\mu_i, \varnothing) \quad (4.2)$$

, where *g()* is an identity or log link function, $E(y_i)$ is the expected value of the response variable $y_i$, $\beta_0$ is the model intercept and $f_j$ is a smooth function of the predictor $x_{ji}$.

The GAMM in Equation 4 can be viewed as a GLMM, where the smoothing parameters are related to variance components (Fahrmeier et al., 2013; Wood, 2017). Hence, assuming any of the above distributions, a GAMM modelling an outcome variable that includes a subject-level random factor ϒ with random intercept and random slopes, and a time series with residual autocorrelation, can be generally expressed as:

$$g(E[y_i]) = \beta_0 + s_1(x_{1i}) + s_2(\gamma_{1i} \cdot x_{1i}) + s_3(x_{2i}) + s_4(\gamma_{1i} \cdot x_{2i}) + s_5(x_{3i}) + s_6(\gamma_{1i} \cdot x_{3i}) + s_7(x_{4i}) + \gamma_{0i} + \text{[illegible]} \quad (5.1)$$

, where $\varepsilon_i$ is the remaining random error term including first-order autocorrelation term AR(1). Substituting by the predictors under study (seasonal depth, amplitude, mean gradient and time) our GAMM model takes the following form:

$$g(E[depth_i]) = \beta_0 + s(sesonal_{depthi}) + s(fishID_i \cdot sesonal_{depthi}) + s(amplitude_i) + s(fishID_i \cdot an \text{[illegible]} \quad (5.2)$$

, where:
$E(y_{ij})$ is the expected value of depth for a fish $id_i$ at time $t_i$,
$\beta_0$ is the population mean depth for a given species and diel period,
$fishID_{0i}$ is the random intercept generating an i.i.d. Gaussian random coefficient for each level of the factor variable *fishID* (different levels for each species), and model matrix component defined by model.matrix(~1:$fishID^{-1}$) that is added to the whole model matrix,
*s()* are smoothing functions (penalized regression splines) of the covariates,
$sesonal\_depth_{1i}$, $amplitude_{2i}$ and $mean\_gradient_{3i}$ are main-effects smoothers related to average fixed effects,

$t_{ij}$ is the average effect of time,
$fishID_{1i} \cdot sesonal\_depth_{1i}$, $fishID_{1i} \cdot amplitude_{2i}$ and $fishID_{1i} \cdot mean\_gradient_{3i}$ are random slopes defining separate i.i.d. Gaussian random slope effects relating depth to each covariate for each level of *fishID*, with model matrices defined by model.matrix($\sim X_{ji}$:$fishID^{-1}$) and added to the whole model matrix,
$\varepsilon_t$ is the residual temporal correlation of the series $\{t = t_1,\ldots,t_n\}$ with $\varepsilon_t = W + u_t$, where $W = \rho\varepsilon_t$ is a covariance matrix for a first-order autocorrelation coefficient $-1<\rho>1$, and $u_t$ is assumed to be i.i.d. Gaussian with mean $E[u_t] = 0$ and variance $\sigma_u^2$ .

##### *1.2. The model in R*

In R code, the model in Equation 5.2 is fit using the bam() function in the *mgcv* package:

```
m_BRNs <- bam(formula = det_depth ~
                s(sesonal_depth, bs = 'ts', k = 10) +
                s(amplitude, bs = 'ts', k = 10) +
                s(mean_gradient, bs = 'ts', k = 10) +
                s(time, bs = 'ts') +
                s(sesonal_depth, fishid, bs = 're', m = 1) +
                s(amplitude, fishid, bs = 're', m = 1) +
                s(mean_gradient, fishid, bs = 're', m = 1) +
                s(fishid, k = 19, bs = 're', m = 1),
                data = data,
                family = 'gaussian', method = 'REML',
                nthreads = 10, cluster = 10, gc.level = 0,
                AR.start = startindex, rho = rho_start_value)
```

*($X_{ji}$, bs='ts')* correspond to average main effects, *s(fishID, bs='re')* is the fish-specific means (random intercepts) and *($X_{ji}$, fishID, bs='re')* the fish-specific effects (random slopes) of the three covariates.

When the model is fit, the maximum number of basis functions that a smooth term actually uses is indicated by k' where always k' < k, and the estimated effective degrees of freedom (EDF) are an indicative of the level of non-linearity of the relationships between predictors and response variable. If EDF-1≈1, the smooth term approaches a linear relationship penalized by the model. EDF can therefore be used to evaluate complexity of the smooth terms in the fitted model. Here, we use default ten basis functions (*k=10*).

To fit smooth functions to predictors we use a special class of shrinkage smoothers based on thin plate regression spline (t.p.r.s.) (specified by *bs='ts'*) that implements the smoothing penalty $\beta^T S_i \beta$ through the identity penalty matrix $S$ so that smooth coefficients are shrunk to 0 (see Equation 3). While this approach allows ruling out heavily penalized smooth terms from the model thus performing automatic variable selection (Marra & Wood, 2011), for simplicity we attain to a general model formula with all predictors included in the model, irrespective of the smoothness parameter and/or the level of linearity (EDF) after penalization by the model.

The argument *m=1* explicitly specifies a squared first derivative penalty term that serves to correct uncertainty from the main-effects smoothers (with second derivative penalty by default), thus reducing concurvity between the two terms (Pedersen et al., 2019).

*AR.start* = *startindex* indicates the starting value of an autocorrelation structure AR(1) and *rho* = *rho_start_value*, the autocorrelation coefficient $\rho$ computed from the starting lag value of a model without the autocorrelation term using the start_value_rho() function in the R package *itsadug* (van Rij et al., 2017).
.
We used restricted maximum likelihood smoothing parameter estimation (*method* = *REML*) that treats random effects similarly as in GLMM likelihood based methods (Wood, 2011). In addition, it allows computing confidence intervals of the variance components from the smoothing parameters with the function gam.vcomp(). Note that *mgcv* only supports simple i.i.d. Gaussian random effects and with the above formula it is only possible to estimate uncorrelated random effects but not correlations between the subject-level random intercepts and slopes (and the slope–intercept covariance?).

##### *1.3. Variance components*

As seen before, fitting a GAMM involves fitting penalized smooth functions with a penalty term, $\beta^T S_i \beta$, which becomes a function of an identity penalty matrix $S$ on the model coefficients $\beta$ through shrinkage of $\beta$ toward zero. This is equivalent to the $\Sigma_\theta$ model matrix component of the random effects i.i.d. (Equation 1) including a diagonal matrix Σ of scaling factors $\{s_1,\ldots,s_n\}$ equal to 1 and determined by the scale parameter ϕ, which is added to the whole model matrix $Z_i$ (Wood, 2008, 2017). Therefore, assuming that each random intercept and random slope in our model is a basis function (Equation 5), the penalty matrix $S$ of the random effects i.i.d. is a diagonal matrix with a constant value of 1 where the penalty term is dependent on the smoothing parameter $\lambda$ of the fitted model. Adding the smoothing parameter to the penalty, $\lambda\beta^T S_i \beta$ (Equation 3) the variance of the random effects b, $\Sigma_\theta$ in Equation 4, is given by dividing the scale parameter by the smoothing parameter, $\phi/\lambda$, where ϕ is equivalent to the residual variance component ($V_e$) in a GLMM. Likewise, the variance of $y_i$, *var, is* equal to the scale parameter ϕ (Equation 4) or $V_e$ in a GLMM (Lin & Zhang, 1999; Fahrmeier et al., 2013; Wood, 2017).

In R code we implement gam.vcomp() in a custom function to calculate these variance components (±95% CI) for each of the smooth terms. The function calculates the variance components directly on the original scale by reversing the rescaling of the penalty matrices (Wood, 2008, 2011):

```
# Create a list containing all final fit models

model_BRNs <- list(mdl_tench_day_BRNs,
                   mdl_tench_night_BRNs,
                   mdl_pike_day_BRNs,
                   mdl_pike_night_BRNs,
                   mdl_wels_day_BRNs,
```

```
                mdl_wels_night_BRNs,
                mdl_rudd_day_BRNs,
                mdl_rudd_night_BRNs)

# Function to calculate variance components on the original scale
# Otherwise if only gam.vcomp() used sd estimates are computed

for(model in model_BRNs){
       vars <- function(model){
       var_rsc <- c(model$reml.scale/(model$sp[1]/model$smooth[[1]]$S.scale),
                 model$reml.scale/(model$sp[2]/model$smooth[[2]]$S.scale),
                 model$reml.scale/(model$sp[3]/model$smooth[[3]]$S.scale),
                 model$reml.scale/(model$sp[4]/model$smooth[[4]]$S.scale),
                 model$reml.scale/(model$sp[5]/model$smooth[[5]]$S.scale),
                 model$reml.scale/(model$sp[6]/model$smooth[[6]]$S.scale),
                 model$reml.scale/(model$sp[7]/model$smooth[[7]]$S.scale),
                 model$reml.scale/(model$sp[8]/model$smooth[[8]]$S.scale),
                 scale = model$reml.scale)
        vars <- data.frame(var = var_rsc, gam.vcomp(model))
               print(vars)
}}
```

##### *1.4. Repeatability estimation*

We analyzed the consistency of fish depths across the range of amplitude values. For that we calculated the repeatability index (*R*) as the proportion of the total phenotypic variance ($V_{total}$) that is due to inter-.individual differences ($V_{fishID}$) (Nakagawa & Schielzeth, 2010). In a mixed-effects model $V_{total}$ is partitioned in two components, inter-individual variance ($V_{fishID}$) and intra-individual variance (i.e., residual variance, $V_e$) which in GAMM are equivalent to the smoothing random intercept $s(\tau_{00})$ and the scale parameter ϕ, respectively (see section above). Thus:

$$R = \frac{V_{fishID}}{V_{fishID} + V_e} = \frac{s(\tau_{00})}{s(\tau_{00}) + \varnothing} \tag{6}$$

For each species and diel period we calculate both agreement and adjusted *R*. Agreement *R* was estimated from models fitted without fixed effects including only the random intercept *fishID* (model_fishid). Adjusted *R* was estimated either from models including the fixed effects of the covariates plus the random intercept (model_fixed) or from models with additional random slopes for each covariate (model_BRNs). In both cases, *R* is assumed to be constant for all values of the confounding (fixed and random) variables. We use the following code to compute R estimates across models and put them in a table (Table 3):

```
# Create additional lists for random-intercept and fixed-effects models

model_fishid <- list(mdl_tench_day_fishid,
                 mdl_tench_night_fishid,
                 mdl_pike_day_fishid,
                 mdl_pike_night_fishid,
```

```
                    mdl_wels_day_fishid,
                    mdl_wels_night_fishid,
                    mdl_rudd_day_fishid,
                    mdl_rudd_night_fishid)

model_fixed <- list(mdl_tench_day_fixed,
                   mdl_tench_night_fixed,
                   mdl_pike_day_fixed,
                   mdl_pike_night_fixed,
                   mdl_wels_day_fixed,
                   mdl_wels_night_fixed,
                   mdl_rudd_day_fixed,
                   mdl_rudd_night_fixed)


# Create a list for each species and diel period

names <- c("Tench-Day", "Tench-Night", "Pike-Day", "Pike-Night", "Wels-Day",
                "Wels-Night", "Rudd-Day", "Rudd-Night")


# Function for calculating unadjusted and adjusted R (± 95% CI) using variance components

for(model in model_fishid){
unadj.R <- function(model){
 vars <- variance_comp(model, rescale = TRUE, coverage = 0.95)
 unadj.R <- (vars$variance)[1]/((vars$variance)[1] + (vars$variance)[2])
 unadj.R_lower_ci <- (vars$lower_ci)[1]^2/((vars$lower_ci)[1]^2 + (vars$lower_ci)[2]^2)
 unadj.R_upper_ci <- (vars$upper_ci)[1]^2/((vars$upper_ci)[1]^2 + (vars$upper_ci)[2]^2)
    print(c(unadj.R_lower_ci,unadj.R,unadj.R_upper_ci))

}}

for(model in model_fixed){
adj.fix.R<- function(model){
 vars <- variance_comp(model, rescale = TRUE, coverage = 0.95)
 adj.fix.R <-(vars$variance)[5]/((vars$variance)[5] + (vars$variance)[6])
 adj.fix.R_lower_ci <- (vars$lower_ci)[5]^2/((vars$lower_ci)[6]^2 + (vars$lower_ci)[6]^2)
 adj.fix.R_upper_ci <- (vars$upper_ci)[5]^2/((vars$upper_ci)[6]^2 + (vars$upper_ci)[6]^2)
    print(c(adj.fix.R_lower_ci,adj.fix.R,adj.fix.R_upper_ci))
}}

for(model in model_BRNs){
adj.BRNs.R <- function(model){
 vars <- variance_comp(model, rescale = TRUE, coverage = 0.95)
 adj.BRNs.R <- (vars$variance)[8]/((vars$variance)[8] + (vars$variance)[9])
 adj.BRNs.R_lower_ci <-(vars$lower_ci)[8]^2/((vars$lower_ci)[8]^2 + (vars$lower_ci)[9]^2)
 adj.BRNs.R_upper_ci <-(vars$upper_ci)[8]^2/((vars$upper_ci)[8]^2 + (vars$upper_ci)[9]^2)
    print(c(adj.BRNs.R_lower_ci,adj.BRNs.R, adj.BRNs.R_upper_ci))
}}


# Create separate tables for unadjusted and adjusted R

table_unadj.R <- lapply(model_fishid, unadj.R)
df_unadj.R <- as.data.frame(data.table::transpose((table_unadj.R)))
colnames(df_unadj.R) <- c("lower CI", "unadj.R", "upper CI")
unadj_R <- zapsmall(df_unadj.R, digits=6)
unadj_R <- cbind(ID = 1:nrow(unadj_R), unadj_R)    # add index applying cbind function
unadj_R$ID <- as.numeric(unadj_R$ID)
```

```
table_adj.fix.R <- lapply(model_fix, adj.fix.R)
df_adj.fix.R <- as.data.frame(data.table::transpose((table_adj.fix.R)))
colnames(df_adj.fix.R) <- c("lower CI", "adj.fix.R", "upper CI")
adj_fix_R <- zapsmall(df_adj.fix.R, digits=6)
adj_fix_R <- cbind(ID = 1:nrow(adj_fix_R), adj_fix_R)
adj_fix_R$ID <- as.numeric(adj_fix_R$ID)

table_adj.BRNs.R <- lapply(model_BRNs, adj.BRNs.R)
df_adj.BRNs.R <- as.data.frame(data.table::transpose((table_adj.BRNs.R)))
colnames(df_adj.BRNs.R) <- c("lower CI", "adj.BRNs.R", "upper CI")
adj_BRNs_R <- zapsmall(df_adj.BRNs.R, digits=6)
adj_BRNs_R <- cbind(ID = 1:nrow(adj_BRNs_R), adj_BRNs_R)
adj_BRNs_R$ID <- as.numeric(adj_BRNs_R$ID)

# Merge the three tables

All_R <- merge(unadj_R, adj_fix_R, by =c("ID"),all=TRUE)
All_R <- merge(All_R, adj_BRNs_R, by =c("ID"),all=TRUE)
rownames(All_R) <- names
```

#### 1.5. *Plotting reaction norms to seiche and individual variability*

```
# Plot individual fish slopes using the exponential transform function for non-gaussian
distributions

plot_smooth(mdl_tench_day_BRNs, view = "amplitude", hide.label = TRUE,
        legend_plot_all = FALSE, plot_all = "fishid", xlab = "Amplitude (seiche
        strength)", ylab = "Depth [m]", main = "Tench - Day", col = cbPalette[1:20])

plot_smooth(mdl_tench_night_BRNs, view = "amplitude", transform=exp, hide.label = TRUE,
        legend_plot_all = FALSE, plot_all = "fishid", xlab = "Amplitude (seiche
        strength)", ylab = "Depth [m]", main = "Tench - Night", col = cbPalette[1:20])

plot_smooth(mdl_pike_day_BRNs, view = "amplitude", hide.label = TRUE, legend_plot_all =
        FALSE, plot_all = "fishid", xlab = "Amplitude (seiche strength)", ylab =
        "Depth [m]", main = "Pike - Day", col = cbPalette[1:20])

plot_smooth(mdl_pike_night_BRNs, view = "amplitude", hide.label = TRUE, legend_plot_all =
        FALSE, plot_all = "fishid", xlab = "Amplitude (seiche strength)", ylab =
        "Depth [m]", main = "Pike - Night", col = cbPalette[1:20])

plot_smooth(mdl_wels_day_BRNs, view = "amplitude", transform=exp, hide.label = TRUE,
        legend_plot_all = FALSE, plot_all = "fishid", xlab = "Amplitude (seiche
        strength)", ylab = "Depth [m]", main = "Wels - Day", col = cbPalette[1:20])

plot_smooth(mdl_wels_night_BRNs, view = "amplitude", transform=exp, hide.label = TRUE, =
        legend_plot_all = FALSE, plot_all = "fishid", xlab = "Amplitude (seiche
        strength)", ylab = "Depth [m]", main = "Wels - Night", col = cbPalette[1:20])

plot_smooth(mdl_rudd_day_BRNs, view = "amplitude", hide.label = TRUE, legend_plot_all =
        FALSE, plot_all = "fishid", xlab = "Amplitude (seiche strength)", ylab =
        "Depth [m]", main = "Rudd - Day", col = cbPalette[1:20])

plot_smooth(mdl_rudd_night_BRNs, view = "amplitude", transform=exp, hide.label = TRUE,
        legend_plot_all = FALSE, plot_all = "fishid", xlab = "Amplitude (seiche
        strength)", ylab = "Depth [m]", main = "Rudd - Night", col = cbPalette[1:20])
```

```
# Using a loop function only (if no transformation is done)

for(i in model_BRNs){
    par(mfrow=c(2,8))
    names <- c("Tench-Day", "Tench-Night", "Pike-Day", "Pike-Night",
               "Wels-Day","Wels-Night", "Rudd-Day", "Rudd-Night")
    plot_smooth(i, view = "amplitude", hide.label = TRUE, legend_plot_all = FALSE,
    plot_all = "fishid", xlab = "Amplitude (seiche strength)", ylab = "Depth [m]",
    main = paste0(" ", names), col = cbPalette[1:20])
}


# Define a list of datasets for each species and diel period

Datasets <- list(data_tench_day_5min,
             data_tench_night_5min,
            data_pike_day_5min,
            data_pike_night_5min,
            data_wels_day_5min,
            data_wels_night_5min,
            data_rudd_day_5min,
            data_rudd_night_5min)


# Function to plot individual variation across amplitude range

for(df in datasets){
plot.var <- function(df){
        mid <- mean(df$amplitude)
        df <- df[complete.cases(df), ]
        df$fishid = reorder(df$fishid, df$det_depth, FUN = mean)
        ggplot(df, aes(fishid, det_depth)) +
        geom_point(aes(color = amplitude_cat), size = 2) +
        theme_bw() + theme(panel.grid.major = element_blank(),
        panel.grid.minor = element_blank(),
        axis.text.x=element_text(angle=90,hjust=1)) +
        scale_color_gradient2(midpoint = mid, low = cbPalette[7],
        mid = "white", high = cbPalette[4], space = "Lab" ) +
        scale_y_reverse() +
        xlab("Fish ID") + ylab("Depth [m]") + labs(colour = "amplitude")
}}

ptd<-plot.var(data_td) + annotate("text", x = 16, y = 0, label=("italic('R') == '0.10'"),
   size = 4, parse = TRUE)
ptn<-plot.var(data_tn) + annotate("text", x = 16, y = 0, label=("italic('R') == '0.19'"),
   size = 4, parse = TRUE)
ppd<-plot.var(data_pd) + annotate("text", x = 16, y = 0, label=("italic('R') == '0.98'"),
   size = 4, parse = TRUE)
ppn<-plot.var(data_pn) + annotate("text", x = 16, y = 0, label=("italic('R') == '0.52'"),
   size = 4, parse = TRUE)
pwd<-plot.var(data_wd) + annotate("text", x = 16, y = 0, label=("italic('R') == '0.40'"),
   size = 4, parse = TRUE)
pwn<-plot.var(data_wn) + annotate("text", x = 16, y = 0, label=("italic('R') == '0.37'"),
   size = 4, parse = TRUE)
prd<-plot.var(data_rd) + annotate("text", x = 16, y = 0, label=("italic('R') == '0.11'"),
   size = 4, parse = TRUE)
prn<-plot.var(data_rn) + annotate("text", x = 16, y = 0, label=("italic('R') == '0.10'"),
   size = 4, parse = TRUE)


# Panel plot
```

```
plot_grid(
ptd + ggtitle("Tench-Day") + theme(plot.title = element_text(hjust = 0.5, color =
     "black", size = 12, face = "bold")),
ptn + ggtitle("Tench-Night") + theme(plot.title = element_text(hjust = 0.5, color =
     "black", size = 12, face = "bold")),
ppd + ggtitle("Pike-Day") + theme(plot.title = element_text(hjust = 0.5, color =
     "black", size = 12, face = "bold")),
ppn + ggtitle("Pike-Night") + theme(plot.title = element_text(hjust = 0.5, color =
     "black", size = 12, face = "bold")),
pwd + ggtitle("Wels-Day") + theme(plot.title = element_text(hjust = 0.5, color =
     "black", size = 12, face = "bold")),
pwn + ggtitle("Wels-Night") + theme(plot.title = element_text(hjust = 0.5, color =
     "black", size = 12, face = "bold")),
prd + ggtitle("Rudd-Day") + theme(plot.title = element_text(hjust = 0.5, color =
     "black", size = 12, face = "bold")),
prn + ggtitle("Rudd-Night") + theme(plot.title = element_text(hjust = 0.5, color =
     "black", size = 12, face = "bold")),
align = "hv",
axis = 'tblr',
label_fontface = "bold",
label_fontfamily = "Times New Roman",
label_size = 8,
rel_heights= c(1,1),
ncol = 4,
nrow = 2,
hjust = -0.7, label_x = 0.01
)
```
